# Substrate rigidity modulates traction forces and stoichiometry of cell matrix adhesions

**DOI:** 10.1101/2021.10.29.466476

**Authors:** Hayri E Balcioglu, Rolf Harkes, Erik HJ Danen, Thomas Schmidt

## Abstract

In cell matrix adhesions, integrin receptors and associated proteins provide a dynamic coupling of the extracellular matrix (ECM) to the cytoskeleton. This allows bidirectional transmission of forces between the ECM and the cytoskeleton, which tunes intracellular signaling cascades that control survival, proliferation, differentiation, and motility. The quantitative relationships between recruitment of distinct cell matrix adhesion proteins and local cellular traction forces are not known. Here, we applied quantitative superresolution microscopy to cell matrix adhesions formed on fibronectin-stamped elastomeric pillars and developed an approach to relate the number of talin, vinculin, paxillin, and focal adhesion kinase (FAK) molecules to the local cellular traction force. We find that FAK recruitment does not show an association with traction-force application whereas a ~60 pN force increase is associated with the recruitment of one talin, two vinculin, and two paxillin molecules on a substrate of effective stiffness of 47 kPa. On a substrate with a four-fold lower effective stiffness the stoichiometry of talin:vinculin:paxillin changes to 2:12:6 for the same ~60 pN traction force. The relative change in force-related vinculin recruitment indicates a stiffness-dependent switch in vinculin function in cell matrix adhesions. Our results reveal a substrate-stiffness-dependent modulation of the relation between cellular traction-force and the molecular stoichiometry of cell-matrix adhesions.

## Introduction

Cell matrix adhesions couple the intracellular cytoskeletal network to the extracellular matrix (ECM) and are key sites for bidirectional mechanotransduction ^1^. First, they are the sites where cells apply myosin-driven contractile forces to their environment, for instance during cell migration and tissue remodeling ^2^. Second, they allow cells to sense and respond to changes in stiffness of their environment. The latter represents an important mechanical cue regulating stem cell differentiation, cancer progression, and various other processes in life and disease ^1,3,4^.

Cell matrix adhesions contain integrin transmembrane receptors that bind ECM components with their globular extracellular head domains. On the cytoplasmic side, integrins connect to a large complex of associated proteins with their intracellular tail domains. Integrins and integrin-associated proteins in cell matrix adhesions have been demonstrated to change conformation thereby exposing new interaction sites when stretched by force ^5^. Several of the associated proteins, including talin and vinculin connect the integrin cytoplasmic tails to the F-actin network ^6^. Others, such as paxillin and focal adhesion kinase (FAK) are involved in local signaling platforms that regulate actin cytoskeletal dynamics for instance through Rho GTPases ^7^. This allows cellmatrix adhesions to adjust their molecular architecture in response to force, ensuring a balance between extracellular (ECM) and intracellular forces.

Cell-matrix adhesions are highly dynamic structures ^8^. Superresolution microscopy techniques have been applied to reveal in detail the 3D multimolecular layered architecture of cell-matrix adhesions ^9,10^. It is well known that larger cell-matrix adhesions support higher forces ^11–13^ but quantitative relationships between recruitment of individual cell-matrix adhesion proteins and local traction-force application have not been reported. Here, we developed a novel method for the analysis of antibody-mediated direct stochastic optical reconstruction microscopy (dSTORM) ^14^ images. For transformation of dSTORM data into local molecule-counting, we followed a real space approach making optimal use of the high positional accuracy characteristic of super-resolution imaging. The method has similarities to the number and brightness analysis known from correlation imaging ^15^ and the Fourier ring-correlation analysis method ^16^. We applied this novel method to four distinct cellmatrix adhesion components, talin, vinculin, paxillin, and FAK. The combination of dSTORM with traction-force microscopy allowed us to unravel quantitative relationships between their recruitment to cell matrix adhesions and local traction forces.

For cells plated on a substrate with an effective Young’s modulus of 47 kPa, we determined that addition of one talin, vinculin, and paxillin molecule to a cell matrix adhesion is accompanied by an on average 66, 30, and 32 pN increase in local traction-force, respectively. On a 12 kPa substrate the stoichiometry for talin:vinculin:paxillin changes from 1:2:2 per ~60 nN force increment to approximately 2:12:6 for the same amount of traction force. Surprisingly, FAK recruitment did not significantly correlate with local traction force increase, irrespective of substrate rigidity. These findings provide a first quantitative relationship between recruitment of distinct cell matrix adhesion proteins and local traction forces and reveals remarkable regulation of the stoichiometry by substrate stiffness.

## RESULTS

### dSTORM on cell matrix adhesion proteins

We analyzed the organization of vinculin in cell matrix adhesions of vinculin knockout mouse embryonic fibroblasts transiently expressing GFP-vinculin by confocal and super-resolution optical microscopy. Comparing signals derived from an Alexa-532-conjugated GFP nanobody to those from an Alexa-647-conjugated secondary antibody targeting a monoclonal vinculin antibody with confocal microscopy showed that these signals co-localized only in GFP positive cell matrix adhesions as expected (Figure 1A, top, red arrows; i,ii). As a control, cell matrix adhesions in vinculin null MEFs lacking GFP-vinculin were readily identified using a paxillin antibody (Figure 1A, bottom, green arrows; iii) while such adhesions did not stain when the vinculin antibody was used (Figure 1A, top).

**Figure 1:**
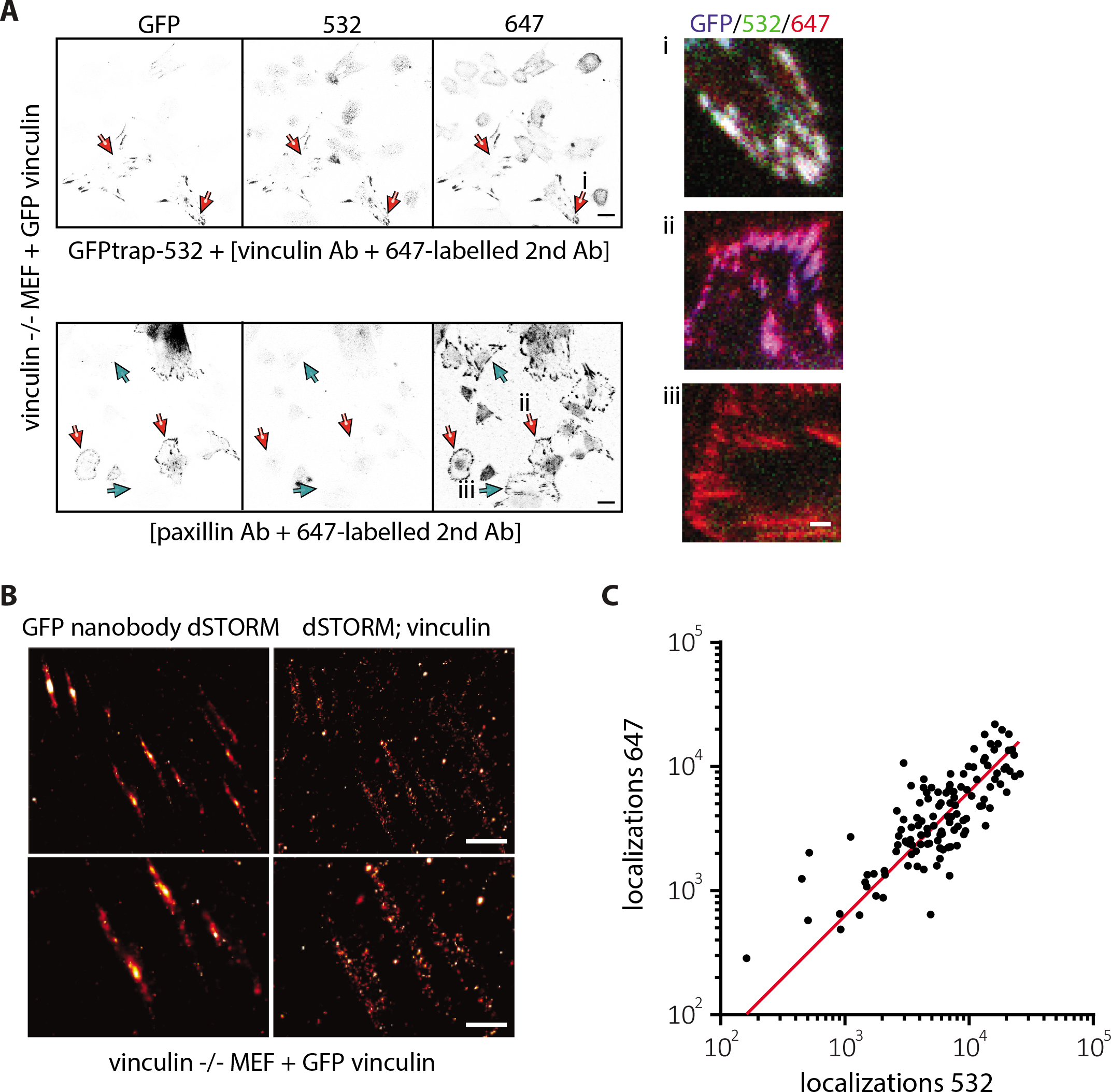
dSTORM on cell matrix adhesions. **A,** confocal images of vinculin -/- MEFs transiently expressing GFP vinculin, immunostained with the indicated antibodies. Red arrows indicate cells that are GFP (and vinculin) positive, green arrows indicate cells that are GFP (and vinculin) negative. In i, ii and iii merged images for zoom-ins of the indicated adhesions are shown. **B,** representative dSTORM images of cells immunostained with GFP nanobody conjugated with Alexa532 obtained with 532 nm laser (left) and [vinculin antibody plus secondary antibody conjugated with Alexa647] obtained with 647 nm laser (right). **C,** comparison of number of localizations obtained from individual adhesions by applying dSTORM to first Alexa647 and then to Alexa532. Red line indicates the linear fit (R^2^=0.47). Scale bars are 20 μm (A, left panels), 3 μm (A, right panels i-iii), 100 nm (B, top) and 50 nm (B, bottom).

Subsequently, we performed dSTORM and analyzed the overlap between fluorophore localizations for Alexa-532-conjugated GFP nanobody signals with vinculin monoclonal antibody followed by Alexa-647-conjugated secondary antibody signals in vinculin null/GFP-vinculin cells. Samples were immersed in a switching-buffer, which lead to quenching of the fluorescence with brief, infrequent de-quenching events resulting in images of sparse signal density < 0.1 μm ^-2^. This permitted detection of signals from individual fluorophores on a sensitive camera (for details see the Material&Methods section). After localization of all fluorophores in each frame, the image of the cell matrix adhesion structure was reconstructed from an image stack of 2×10^4^ frames. dSTORM images, similar to confocal images, showed the predicted overlap between Alexa-532 and Alexa-647 localizations (Figure 1B). The number of localizations obtained from the two different fluorophores across 105 adhesions in 11 different cells as determined by dSTORM followed a linear dependence (Figure 1C).

### Combination of dSTORM and cellular traction force measurements

In order to combine force measurements and super resolution microscopy, we seeded NIH3T3 fibroblasts on fibronectin stamped elastomeric micropillars. On these substrates talin staining followed by confocal imaging identified cell matrix adhesions coupled to fibronectin-stamped μ-pillars (Figure S1A). The average background deflection, corresponding to the displacement resolution, was 50±20 nm as determined by epi-fluorescence imaging in a cell-free region in the field of view of dSTORM imaging (Figure S1B,C). For μ-pillar arrays with effective Young’s moduli of 12 kPa or 47 kPa (spring constants per pillar of 16 nN/μm or 66 nN/μm, respectively), the displacement resolution of 50 nm corresponded to a force precision of 0.8 nN and 3 nN, respectively. Combining epi-fluorescence pillar displacements and dSTORM, provided simultaneous visualization of traction force and localizations in cell matrix adhesions (Figure 2A). The significant increase in spatial resolution between the dSTORM image and a wide-field image for a talin staining is apparent from Figure 2A. The resolution in the dSTORM image was largely determined by the positional accuracy by which each of the molecules was localized. We set the imaging conditions such that on average a signal of 520 photons per localization was detected (Figure 2B,C). At such count rate the localization precision was 14±5 nm (Figure 2B,C).

**Figure 2:**
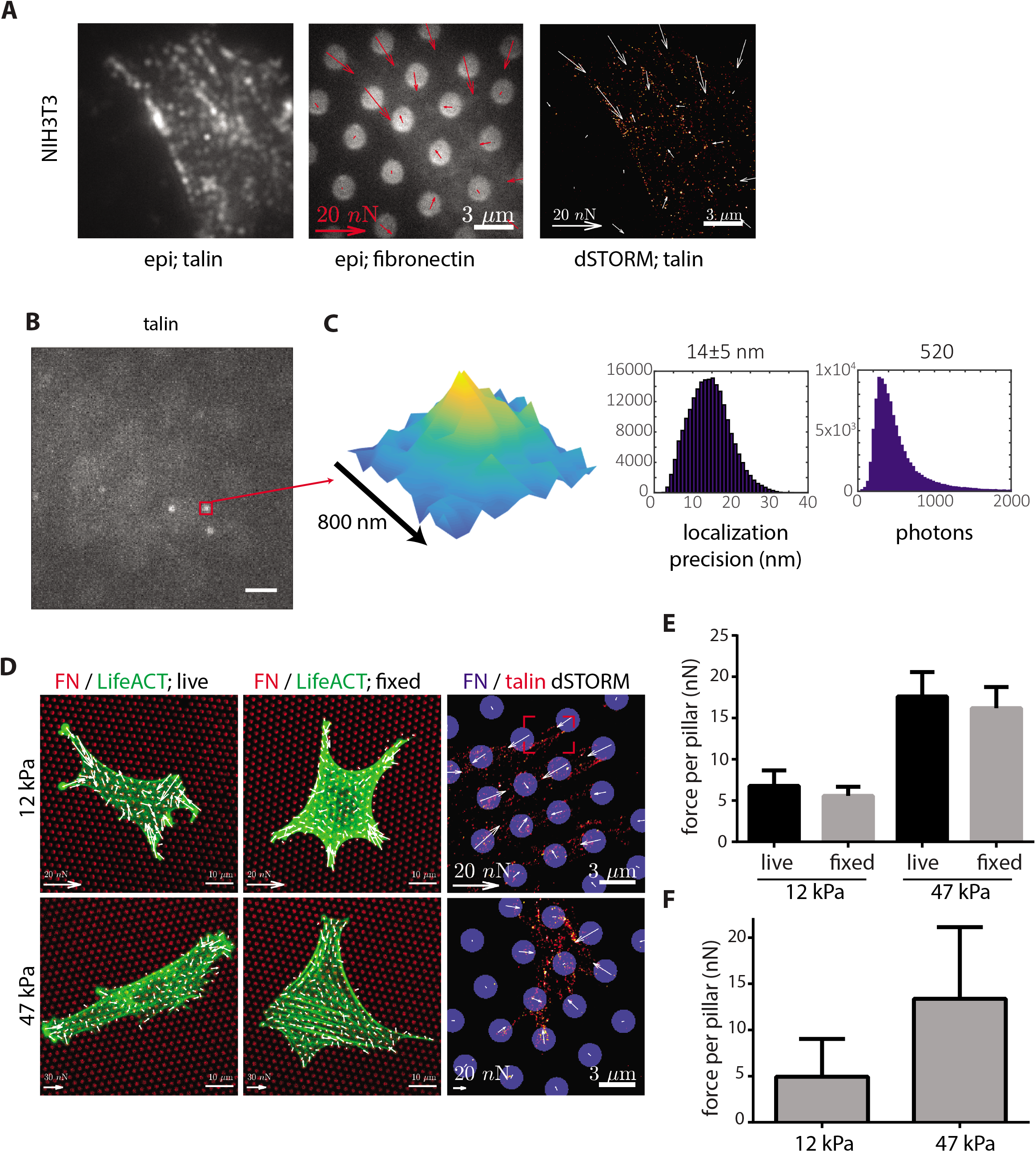
Combination of dSTORM with micropillars. **A,** NIH3T3 cell imaged with the dSTORM setup using epi-fluorescence with 647 (left) and 405 (middle) or dSTORM with 647 channel (right) together with accompanying force measurements (arrows in middle and right image). **B,** example image frame from dSTORM acquisition with several Alexa647 molecules fluorescing. **C,** zoom in of the red square in B (left) and histograms showing the positional accuracy of localizations from this dSTORM acquisition (left) and the intensity of localizations (right).**D,** images of live (left) and fixed (middle and right) NIH3T3 cells on pillars of effective Young’s modulus of 12 kPa (top) or 47 kPa (bottom) stamped with fibronectin conjugated to Alexa647 (left and middle) or to Alexa405 (right). mCherry-LifeACT (left and middle) or talin immunostaining (right; secondary antibody conjugated with Alexa647) was imaged using confocal imaging (left and middle) or dSTORM setup (right) with calculated forces (arrows). **E,F**, bar graphs showing mean ± standard deviation of cellular forces applied per pillar calculated from confocal images (E) or images obtained with dSTORM setup (F) for cells on pillars with indicated stiffnesses. Scale bars are 3 μm (A, D-right), 2 μm (B), 10 μm (D, left and middle); deflection arrow scales are 20 nN (A, D, top-right and bottom) and 30 nN (D, top-left and top-middle).

In order to confirm the force measurements in combination with dSTORM, we compared the measured forces to those obtained from live-cell experiments. We established that forces measured in samples fixed for dSTORM application were slightly lower than forces measured by live confocal imaging of pillar deflections in mCherry-lifeact-labelled NIH3T3 cells (Figure 2D,E). The increase in force, induced by seeding cells on a substrate with higher effective Young’s modulus as measured postfixation recapitulated the increase measured in live cells, as established previously for fixation for standard immunofluorescence ^17^. In accordance with the results obtained by confocal imaging, forces determined by epi-fluorescence microscopy in the field of view of dSTORM imaging of NIH3T3 cells that were immunostained for talin showed a ~3-fold increase for cells seeded on 47 kPa as compared to forces applied by cells seeded on 12 kPa (Figure 2D-F).

### From dSTORM localizations to molecule counts

We next investigated methods for obtaining molecule counts from dSTORM images. For photo activation localization microscopy (PALM) where each protein of interest is fused to one fluorescent protein that bleaches fast and has very short dark times ^18^, a cleaning algorithm can be used^6^. This removes additional detections from the same fluorophore within the positional accuracy for single molecule detection for a given time interval that safely exceeds the photobleaching time of the fluorophores. This method is susceptible to errors if the density of molecules exceeds the positional accuracy as may occur within cell matrix adhesions, and when multiple fluorophores per protein of interest are present as applies to dSTORM methods. Instead, for quantitative analysis of dSTORM data, density-based clustering algorithms are typically used where detections are categorized based on their local density as members of clusters ^19^. Yet, also those methodologies are susceptible to errors due to local clustering of the proteins.

Here, we performed dSTORM mediated by 2-step antibody staining with Alexa647 fluorophores on the secondary antibody that undergoes stochastic blinking ^14,20,21^. In this design, signal amplification of the signal as of binding of multiple secondary antibodies that all contain multiple fluorophores, will have to be accounted for in order to accurately estimate the true local stoichiometry. Therefore, we developed a method that makes use of the inherent high localization precision and signal amplification present in our setup. We based our methodology on analysis of the inter-localization distance distribution in the images, which in turn was used to distinguish between spatially correlated and uncorrelated localizations. Our novel method makes use of the fact that the statistics associated with fluorescence labeling and photophysics, although partly unknown, are equivalent for all spatially correlated, and spatially uncorrelated localizations.

In order to estimate the number of talin molecules in an adhesion coupled to one pillar (Figure 2D, red box; Figure 3A,B), we determined the statistics for the cumulative distribution function (cdf) of inter-localization distances within the adhesion as shown in Figure 3B. For each distance, r, between 2 localizations, the number of distances smaller than r was determined as a function of the squared distance, r^2^. For a spatially random distribution where the distances are uncorrelated, the distribution function increases linearly with r^2^. In our data, the relationship between the cdf of inter-localization distances and r^2^ displayed two-regimes. A linear regime was observed for large r^2^, reflecting localizations belonging to uncorrelated, hence different talin molecules (Figure 3C). Subtracting this linear relationship from the data, the nonlinear regime remained for small r^2^ (Figure 3D). The non-linear initial increase reflects correlated detections, hence belonging to a single talin molecule or to a cluster of talin molecules. The non-linear regime was subsequently fit to a double exponential:

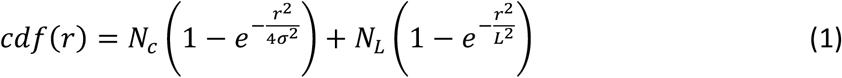

**Figure 3:**
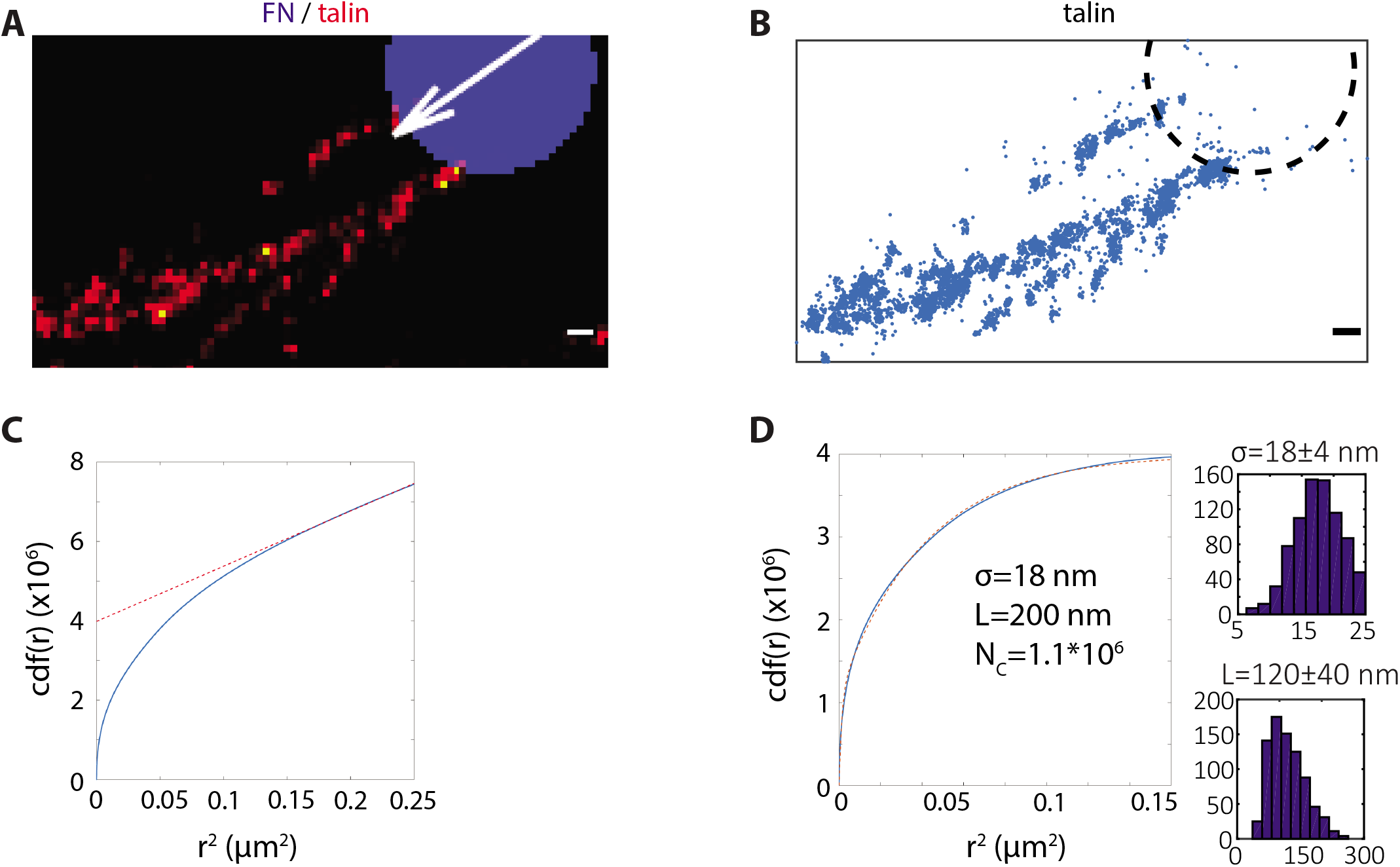
Distribution analysis of talin dSTORM localizations in a single adhesion. **A,** zoom in of the red square from Figure 2D. **B,** image derived from A, showing 6700 localizations of Alexa647 targeted to talin associated with one pillar (dashed circle). **C,** cumulative distance function (cdf) of the localizations in B with a linear line fit (red dashed line) from 0.16 μm^2^ to 0.25 μm^2^. **D,** cdf from C, with linear fit subtracted and accompanying double exponential fit 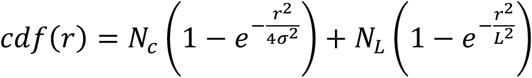 with σ=18 nm, L=200 nm, N_c_=1.10×10^6^ and N_L_=2.9×10^6^. Histograms of fit parameters σ (right-top) and L (right-bottom) obtained across all experiments are shown. Scale bars are 250nm (A, B).

In eq.(1) r denotes the distance between fluorophores, σ the positional accuracy of position detections coupled to a single talin molecule, with N_c_ the corresponding number of correlated distances. Further we added a second exponential that contains the information of the typical size of the focal adhesion, characterized by a typical length-scale L, and the number of localizations within this domain N_L_, which is proportional to the number of talin molecules in close proximity. The structural length scale, L = (120±40) nm, typical for adhesions, was considerably larger than the positional accuracy of talin localizations in dSTORM, σ = (18±4) nm, indicating the two components of the exponential were well separable. From the fit the number of talin molecules Nm in the adhesion (Figure 3D) was the determined by (see materials and methods):

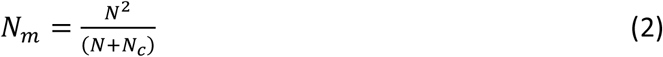

where N is the total number of detections in the image. Our experimental strategy we validated in simulations. We found an excellent agreement between the estimated number of molecules and our analysis: the accuracy was >90% even in conditions of high spatial overlap between individual signals (Supplementary Figure 1D-F). Using our methodology, 40 talin molecules were identified for the adhesion shown in Figure 3A,B.

### Relating the stoichiometry of cell matrix adhesion proteins to traction forces

Having established a methodology to quantify the number of molecules in an adhesion, we studied the stoichiometry of four different cell matrix adhesion proteins in cells seeded on pillar arrays of two different effective Young’s moduli. In the analysis cell matrix adhesion areas were selected and subsequently the corresponding number of talin, vinculin, paxillin, and FAK molecules calculated (see supplementary Figure 2 for histograms for all measurements and calculations). In order to relate the cell’s force application to the number of adhesion molecules, we plotted the local traction force in relation to the number of detected molecules in an adhesion. In total >100 cell-matrix adhesions from 30 NIH3T3 cells was analyzed in 3 independent experiments on μ-pillar arrays of effective Young’s modulus of 47 kPa. The data showed that the cellular force was highly correlated to the number of talin molecules in the respective adhesion (Figure 4A, Supplementary Figure 3). Further, the number of molecules and the force both increased with cell matrix adhesion area (Supplementary Figure 3). Likewise, experiments performed on μ-pillar arrays of lower effective Young’s modulus (12 kPa) showed correlated talin-force relations, yet forces applied by the adhesion per talin molecule were generally lower (Figure 4B, Supplementary Figure 3).

**Figure 4:**
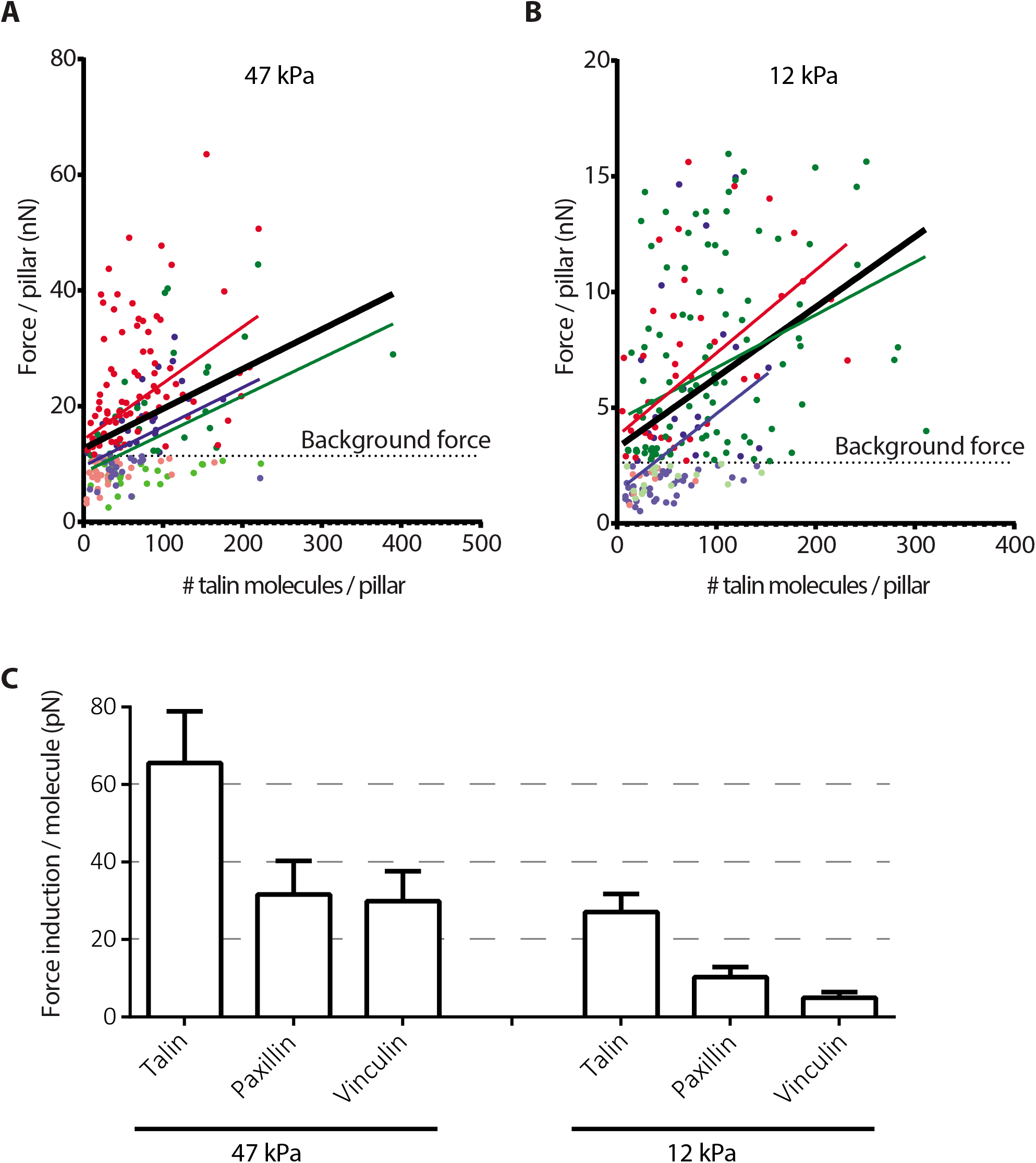
Recruitment of talin, vinculin, and paxillin to cell matrix adhesions is associated with distinct increments in force that depend on substrate stiffness. **A, B,** measured force per cell matrix adhesion plotted against calculated number of talin molecules per adhesion for cells seeded on substrates with effective Young’s modulus 47 kPa (A) and 12 kPa (B). Dots indicate individual adhesions; lines indicate linear fits. Red, green and blue colors represent data from three independent experiments. Solid black line represents linear fit for all data points from all three experiments. Dashed horizontal black line denotes the background forces measured. **C,** bar graphs showing linear fit slope values for relation between local traction force and number of talin, paxillin and vinculin molecules for cells seeded on substrates with effective Young’s modulus 47 kPa and 12 kPa.

Similar to talin, the number of vinculin and paxillin molecules in a cell matrix adhesion correlated with the cellular force application on both 47 kPa and 12 kPa substrates (Supplementary Figure 3). By contrast and to a surprise, the number of FAK molecules in an adhesion was not correlated to the force application on neither of the substrates (Supplementary Figure 3). For FAK the number of molecules per area was uncorrelated to the traction force, indicating that the increased adhesion size rather than an increase in protein density was responsible for the increase in talin, vinculin and paxillin molecules with increased force. For instance, irrespective of the amount of traction force measured, the numbers of talin molecules per square micrometer on 47 kPa and 12 kPa pillars were 52±3 and 54±3, respectively.

Next, we quantified the correlations as linear relation between the abundance of talin, vinculin, and paxillin molecules in a cell matrix adhesion to the local force application. On a substrate of effective Young’s modulus of 47 kPa, for each additional talin molecule an increase in the traction force by 66 pN was determined (Figure 4C). Vinculin and paxillin molecules were associated about half of this force: for each additional vinculin and paxillin molecule, an increase in force by 30 pN and 32 pN was determined, respectively (Figure 4C). On a substrate of effective Young’s modulus of 12 kPa, the force increments were less steep, with a stronger decrease in force association with vinculin was observed: 27 pN/talin, 4.9 pN/vinculin, and 10 pN/paxillin (Figure 4C).

Together our findings indicate that (i) talin, vinculin, and paxillin recruitment to cell matrix adhesions is associated with distinct force increments; (ii) on a substrate of 47 kPa an increase in local traction force of ~60 pN is accompanied by the recruitment of 1:2:2 talin:vinculin:paxillin molecules; (iii) on a four-times softer substrate the force increments per molecule are less pronounced and vinculin-related force decreases dramatically; (iv) FAK recruitment is not significantly associated with the amount of local traction force.

## DISCUSSION

Cell matrix adhesions are highly dynamic multiprotein complexes that allow cells to sense and respond to physical cues from their surrounding ECM. We combined micropillar based traction force microscopy with super resolution microscopy to obtain quantitative relationships between cell matrix adhesion composition and local traction force. In order to address the number of molecules, we used distribution of inter-localization distances in dSTORM images. Our methodology makes use of the high localization precision and signal amplification inherent to dSTORM imaging to quantify the number of molecules.

Previously, a probability density function calculated using Fourier space has been used to correct for multiple detections in PALM (PC-PALM) ^40^. Another method made use of Fourier ring correlation to relate the photophysics of detections outside of the structure of interest with the number of molecules inside this structure ^41^. For either of these techniques to work, the labeling of a protein of interest with a single fluorophore is essential. This requires either precise genome editing or very accurate nanobody labeling. Both of these methods suffer from information loss due to discretization in Fourier space. By contrast, the method we developed here is real space based, taking full advantage of high localization precision. A very different approach was taken where photon arrival times were determined and the antibunching was analyzed to estimate the number of fluorophores present in a focal volume ^42,43^. However, in contrast to our method, those approaches reliy heavily on the detailed knowledge of the photo-physics and so far less been used to count molecules.

Instead of trying to reach a one fluorophore-per-molecule labeling, our methodology actually makes use of the high abundance of fluorophores per protein of interest and combines it with the high localization precision inherent to super resolution methods. Our approach can be readily applied to commercially available antibodies in combination with labeled secondary antibodies used for standard immunostaining. Using this method we reported here distinct force relationships for the cell-matrix adhesion proteins talin, vinculin, and paxillin that are modulated by environmental stiffness, whereas recruitment of downstream FAK molecules was not related to the amount of local traction force (Figure 5).

**Figure 5:**
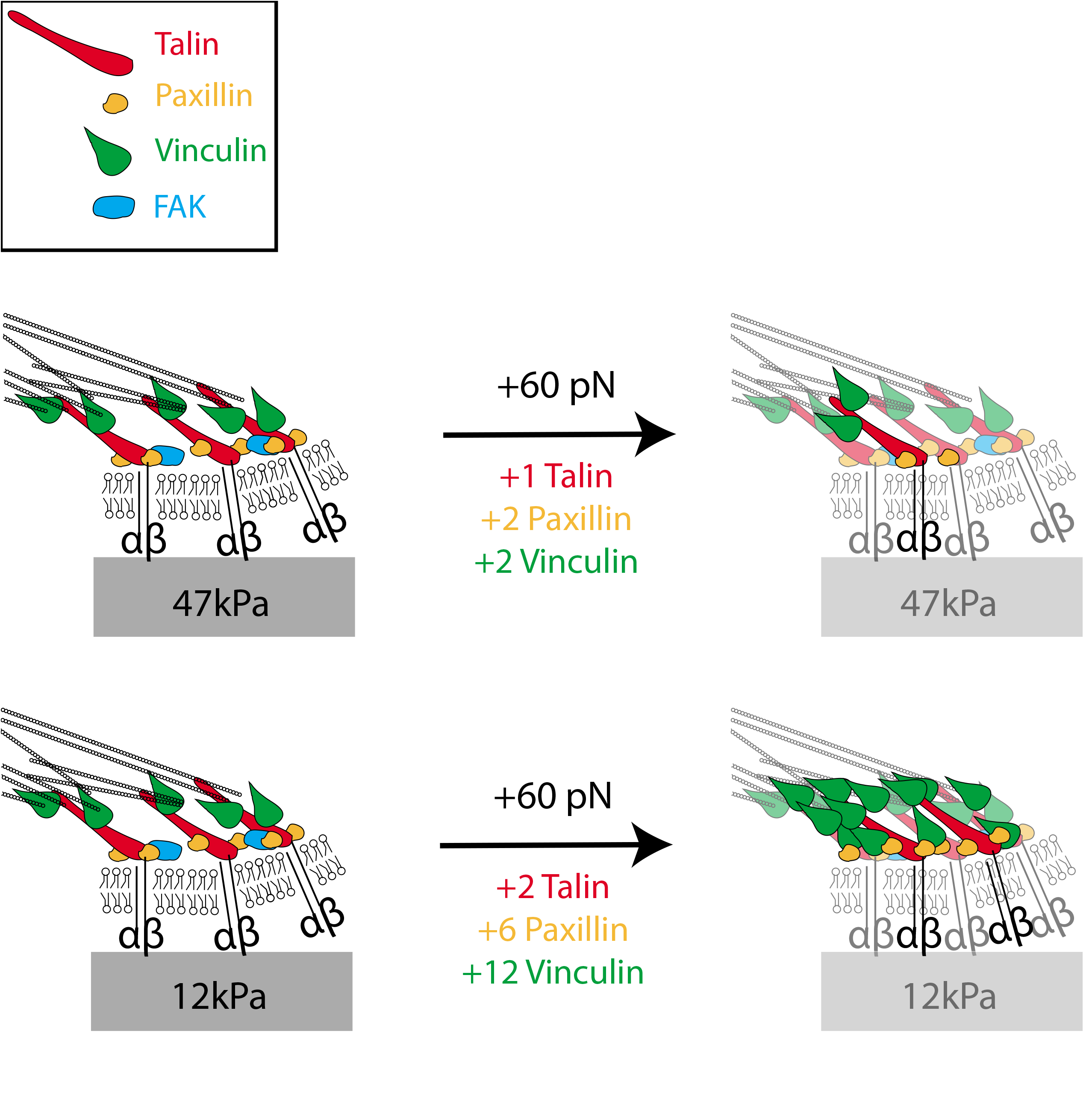
Molecular composition of force responsive cell matrix adhesions. Cartoon depicting recruitment of talin, paxillin and vinculin molecules associated with a ~60 pN increase in force on stiff and soft substrates with indicated effective Young’s moduli.

Similar to previous findings ^11–13^, we observed that larger cell matrix adhesions support higher traction forces. Additionally, we found that recruitment of talin, vinculin, or paxillin to the adhesion with increasing traction force was not accompanied by a significant change in molecule density. Rather, the increase in traction force was supported by the recruitment of cell matrix adhesion proteins causing the adhesion to spatially expand. Recently, it has been reported that tension across individual talin molecules is largely constant over a wide range of substrate rigidities that trigger a range of cellular traction forces ^22^. Together, these findings indicate that an increase in cellular traction force involves expansion of the adhesion through recruitment of new talin molecules without increasing the concentration of talin molecules or the force mediated by individual talin molecules.

Important to note, that the increase in force associated with an additional talin, vinculin, or paxillin molecule that we report here does not represent the actual force exerted on these molecules, but solely represents the overall force on the adhesion. Several additional cell matrix adhesion proteins, that are not analyzed in our current study, are likely to be recruited to the growing adhesion as it applies more traction force. These include proteins that couple integrins to the cytoskeleton, such as α-actinin and filamin ^23^. Their contribution has not been addressed here. As a single talin molecule interacts with a single integrin molecule, our finding that one additional talin molecule is associated with an additional 60 pN traction force on a 47 kPa substrate may point to an additional integrin being recruited to the adhesion with this increase in force. Alternatively, one pre-existing integrin may switch from interaction with filamin or α-actinin to interaction with the newly recruited talin molecule, as an additional 60 pN traction force is applied by the adhesion.

It has been reported that FAK is necessary for cellular traction force generation ^24^ and blocking myosin II activity impairs FAK recruitment to cell matrix adhesions ^25^. In that latter study myosin activity was blocked with blebbistatin (20 μM) resulting in very small adhesions (0.17 μm^2^). Here we showed that FAK recruitment to cell matrix adhesions does not correlate with increased traction forces on short or long pillars. This does not imply that FAK is not implicated in force generation. It has been shown that FAK activation through phosphorylation is force dependent ^26^ and, in turn influences force dependent phosphorylation of paxillin and recruitment of vinculin ^25^. Our findings, together with studies showing that FAK residence times are low and increase with increasing cell matrix adhesion size ^25,27^ suggest that changes in FAK activity rather than its recruitment are coupled to force.

We find that the stoichiometry of talin molecules in cell matrix adhesions is associated with the highest traction forces on stiff as well as softer substrates as compared to vinculin and paxillin. On a stiff substrate an increase in traction force of ~60 pN is associated with one additional talin molecule, whereas two additional vinculin and paxillin molecules are associated with the same force increase (Figure 5). Talin connects the integrin to actin and acts as a scaffold for vinculin binding ^22,28^. Binding of talin to the integrin cytoplasmic tail activates the integrin and enhances ECM binding, and interaction of talin with integrin αvβ3 is important for adhesion strengthening ^29^. Forces on talin molecules open cryptic binding sites for vinculin ^30^. It has been shown that induction of myosin contractility triggers this unfolding, which is also correlated with more actin proximal localization of vinculin and adhesion maturation ^10^. Furthermore, forces transduced by individual talin molecules are reduced in the absence of vinculin but not entirely lost ^22^. These findings indicate that recruitment of talin and vinculin as well as their interaction is important for force related adhesion maturation. As described above, vinculin can also be recruited to cell matrix adhesions through FAK-mediated phosphorylation of paxillin, a process that depends on myosin-mediated contractility ^10,25^

Experiments on isolated talin molecules have shown that cryptic vinculin binding sites become available when talin is under 5-25 pN tension ^31^. The 66 pN or 27 pN increase in traction force we measure for a cell matrix adhesion on a stiff or soft substrate, respectively, per additional talin molecule is beyond the reported threshold for opening vinculin binding sites and below the 100 pN forces that can be supported by single actin molecules ^32^. Notably, vinculin molecules that are recruited to the adhesion via talin, phospho-paxillin or other interactions such as force dependent p130Cas-vinculin binding ^33^, may partially remain in an inactive confirmation, especially on a soft substrate, which may explain the lower force induction measured for each recruited vinculin molecule as compared to talin.

Interestingly, it has been reported that as adhesions enlarge, forces on individual vinculin molecules decrease ^34^. A recent publication shed more light on this by demonstrating a switch behavior for vinculin: for very small and very large adhesions tension on vinculin adhesion growth is slowed, while for adhesions of intermediate size, a positive correlation of vinculin tension with adhesion growth was found ^35^. The fact that vinculin exhibits a slow turnover in cell-matrix adhesions on glass and that inhibition of myosin contractility raises its turnover to that observed for other cell matrix adhesion proteins further suggests that vinculin changes its function with force ^25,36^. Our findings extend these observations, showing that a decrease in substrate rigidity leads to a major decrease in vinculin-associated force: i.e. for the same amount of force increase the number of recruited vinculin molecules is six times higher on a soft versus a stiff substrate (Figure 5). This suggests that the activation of vinculin molecules is less efficient on a low rigidity substrate, rendering a pool of vinculin molecules in an inactive state. Vinculin activation is proposed to occur through its interaction with talin ^10^. Larger forces applied on stiff substrates in our experiments may enhance the talin-vinculin interaction, thereby more effectively supporting vinculin activation and subsequent coupling of vinculin to the actin cytoskeleton.

Taken together, we have combined dSTORM and traction force microscopy to obtain quantitative information on the relationship between the molecular composition of cell matrix adhesions and their force application. We report that an increase in force of ~60 pN is accompanied by recruitment of 1:2:2 talin:vinculin:paxillin proteins on a substrate with an effective stiffness of 47 kPa (Figure 5). The stoichiometry changes on softer substrates, in particular due to a strong reduction of vinculin-associated force. The methodology we introduced here for extraction of quantitative molecular information from super resolution images is readily applicable to other cellular structures given that there is enough signal amplification, i.e. multiple fluorophores associate with the protein of interest and/or multiple blinking events are observed per fluorophore.

## MATERIALS AND METHODS

### Cell culture and transduction

Vinculin KO MEFs (kindly provided by Dr. Johan de Rooij, Utrecht University, NL) and NIH-3T3 fibroblasts were cultured in medium (DMEM; Dulbecco’s modified Eagle’s Medium, Invitrogen/Fisher Scientific) supplemented with 10% new born calf serum, 25 U/ml penicillin and 25 μg/ml streptomycin (Invitrogen/Fisher Scientific cat. # 15070-063). Vinculin KO MEFs were transduced with a GFP-vinculin retroviral construct as previously described ^37^.

### Micropillar preparation and cell seeding

Micropillar arrays were used for cellular traction force measurements according to methodology described previously ^13^. A negative silicon master was made with 10×10 mm arrays of circular holes of 4.1 or 6.9 μm depth, 2 μm diameter and 4 μm center-to-center distance in a hexagonal grid with two rectangular spacers of 10×2 mm wide and 50 μm high aligned on the sides of the arrays using a two-step Deep Reactive Ion Etching (DRIE) process. The negative silicon master was passivated with trichloro silane (Sigma Aldrich) and well-mixed PDMS at 1:10 ratio (crosslinker:prepolymer) was poured over the wafer and cured for 20 hours at 110°C. Pillar arrays of 4.1 and 6.9 μm height had a bending stiffness of 66 nN/μm and 16 nN/μm respectively, corresponding to an effective Young’s modulus, Eeff, of 47.2 and 11.6 kPa respectively ^17^. Stamping of fibronectin was performed using a flat piece of PDMS (1:30 ratio, cured 16 hours at 65°C) that had been incubated with a mix of 50 μg/mL unlabeled fibronectin (Sigma Aldrich) and 10 μg/mL Alexa405 or Alexa647 (both from Invitrogen)-conjugated fibronectin. Subsequently, the stamped micropillars were blocked with 0.2% Pluronic (F-127, Sigma Aldrich) and cells were seeded at single cell density in complete medium and incubated for 5 hours at 37°C and 5% CO_2_.

### Fixation and immunostaining

Samples were washed once with cytoskeleton buffer (CB) (10 mM MES, 150 mM NaCl, 5 mM EGTA, 5 mM MgCl_2_, and 5 mM glucose) ^38^, fixed and permeabilized for 10 seconds with 0.1-0.25% Triton-X, 0.4% paraformaldehyde and 1 μg/mL phalloidin in CB, fixed for 10 min with 4% formaldehyde in CB, permeabilized for 10 min with 0.5% Triton-X, and blocked for 30 min with 0.5% BSA. Immunostaining was performed with an Alexa-532-conjugated GFP nanobody (Chromotek, Germany) or with a primary mouse monoclonal antibody against talin (Sigma, T-3287), FAK (BD Transduction, 610087), paxillin (BD Transduction, 610052) or vinculin (Sigma, V-9131), followed by an Alexa647 conjugated secondary antibody against mouse IgG (Jackson, 115-605-006) following the protocol suggested by Linde et al. ^14^.

### Imaging and analysis

#### dSTORM imaging

Super-resolution imaging was performed on a home-built wide-field single-molecule setup, based on an Axiovert S100 (Zeiss) inverted microscope equipped with a 100x 1.4NA oil-immersion objective (Zeiss, Germany). Micropillar arrays were inverted onto #0 thickness, 25 mm diameter, round coverslips (Menzel Glaser). Imaging was performed in 100 mM mercaptoethylamine (MEA, Sigma Aldrich) in PBS. A 405 nm laser (CrystaLaser, USA) was used for imaging the pillars and photoswitching of the Alexa647 dye to adjust the density of visible fluorophores. The light was reflected into the objective by a dichroic mirror (ZT405/532/635rpc, Chroma, USA). The fluorescence light in the detection path was filtered using the emission filter ZET532/633m (Chroma, USA). Conversion intensities were set between 0 and 250 W/cm^2^ at 405 nm, and the excitation intensity was set to 5 kW/cm^2^ at 647 nm. For each sample, we acquired 20000 images with an acquisition time of 10 ms per frame at a frame rate of 69 Hz. The signal of individual dye molecules was captured on a sCMOS Orca Flash 4.0V2 camera (Hamamatsu, Japan). The average integrated signal of a single fluorophore was 520 detected photons, spatially distributed by the pointspread-function of the microscope of 440 nm FWHM, resulting in a sigma of 187 nm in a Gaussian approximation.

The signal from individual fluorophores was fitted with a 2-dimensional Gaussian using a custom least-squares algorithm in Matlab ^39^. From the fit we determined the location of each fluorophore to an accuracy of 14±5 nm (Fig. 3B). This localization accuracy is slightly higher than its theoretical minimum predicted from the width of the point-spread-function and the detected signal, 187 nm / √[520] = 8.2 nm.

#### Obtaining and fitting the cumulative distribution function

The area of interest from the epi-fluoresence image was obtained by manually drawing a mask containing the cell matrix adhesion extending from the pillar of interest in the direction of the deflection. Localizations corresponding to blinking events in this area were analyzed (Figure 3A,B). From the position data, the two-point spatial correlation function g(r) and subsequently the cumulative distance function (cdf) was calculated

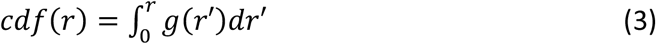

For discrete 2D position data r_i_ = {x_i_, y_i_}, as obtained from the fits, the cdf was constructed as

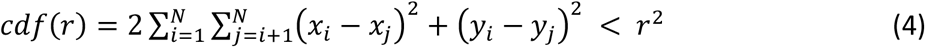

The cdf describes the number of distances that are smaller than r, in a sample of N localizations.

The positional accuracy leads to an apparent Gaussian spread in the localization of an individual molecule characterized by a cdf,

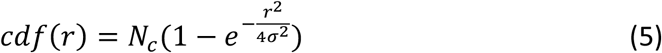

with correlation length, σ_i_, given by a combination of the localization uncertainty for an individual fluorophore and the size of a primary and secondary antibody complex used to label the protein of interest. Both detection and labeling originate from statistical processes. Here N_c_ is the total number of correlated distances and σ the mean positional uncertainty for all localizations coupled to one protein of interest. Eq.(5) is valid for r ~ σ.

On length scales longer than the correlation length the cdf was characterized by a distance distribution for uncorrelated localizations. Assuming a homogeneous, random organization of localizations within a given field-of-view of area, A, the cdf follows a quadratic dependence on distance as:

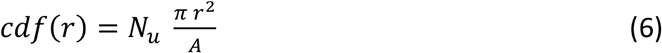

where N_u_ is the number of uncorrelated distances.

Thus, the general form for the spatial correlation function was a linear combination of the correlated and the uncorrelated part:

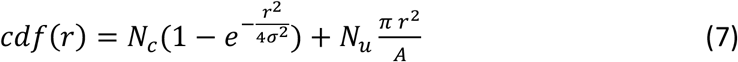

Running a simulation on 2048 individual molecules randomly positioned in a box of 2×2 μm^2^, each detected by 100 localizations at a positional accuracy of 20 nm (Supplementary Figure 1D) the distance distribution was calculated and its dependence on the squared distance, r^2^, is shown (Supplemental Figure 1E). For squared distances beyond 4×10^-3^ μm^2^ the cdf(r) became linearly dependent on r^2^ with the slope of πN_u_/A and y-intersect at N_c_(*2×10^7^*) as predicted.

#### Calculation of number of molecules from the fit

From N_c_ the number of molecules was calculated as follows. The number of localizations, N, originating from N_m_ molecules each being observed n_i_ times is given by

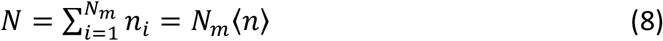

where <n> is the average number of observations per molecule. Hence *N*^2^ = (*N_m_* 〈*n*〉)^2^.

Likewise, the total number of correlated distances, N_c_ per molecule is given by n_i_ × (n_i_-1). For all molecules this yields:

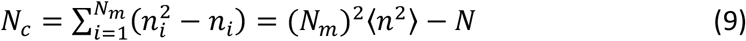

And therefore

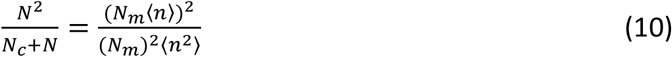

Rearranging gives

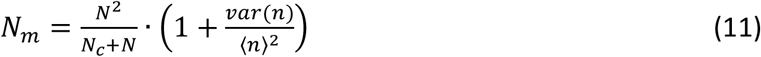

where var(n) = <n^2^>-<n>^2^, is the variance in the number of detections per molecule.

The second term, 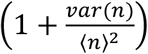, summarizes the properties of the joined statistics of labeling and photophysics of the fluorophores and its value varies between 1 and 2 depending on which of the various processes dominates the joined statistics and for a typical dSTORM experiment is close to one (see the supplementary material for a more detailed analysis).

Simulations were performed for densities between 40 and 4000 randomly distributed molecules on an area of 2×2 μm^2^. One hundred localizations per molecule were simulated with a mean positional accuracy σ = 20 nm. At high densities there was significant overlap of molecules within the image (Supplemental Figure 1D). The number of estimated molecules faithfully followed the input within an error of <10% (Supplemental Figure 1F)

#### Estimation of number of molecules in an adhesion

In the quantification of the number of correlated distances it was assumed that all molecules were randomly organized, which is not the case for molecule clusters such as present in a cell matrix adhesion. This restriction is readily lifted by the addition of a second exponential term with weight, N_L_ that accounts for a length scale, L that characterizes any spatial structures in real data. In this case the cdf for a nonlinear regime becomes

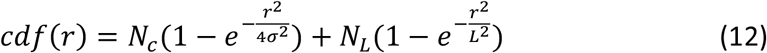

For the distinction of the two components, the typical structural length scale should be significantly larger than the positional accuracy, L > 4σ, typically 40 nm for a positional accuracy of 10 nm. The structure parameter of adhesion clustering measured here was way above this threshold, L = 120±40 nm. Hence the method described above provides a very general solution for molecule counting in superresolution microscopy where

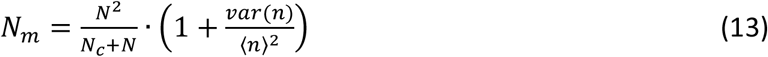

#### Deflection analysis

Pillar deflections were determined at approximately 50 nm precision using a specifically designed Matlab script. The pillar locations were determined from the labeled fibronectin fluorescence images using a fit to the cross-correlation function between a perfect binary circle and the local fluorescence of one pillar. Those positions were compared to those of a perfect hexagonal grid used as reference. From an undeflected array image the accuracy was found to be 47.1 nm (Supplemental Figure 1B,C), which corresponds to a force accuracy of 780 pN and 3.1 nN on the pillar array of Eeff = 11.6 kPa and 47.2 kPa, respectively. Masks for adhesions corresponding to individual pillars of interest were manually drawn for each case.

### Statistical analysis

p-values were calculated using F-test for linear regression analysis using GraphPad Prism 6.0.

## ACKNOWLEDGEMENTS

We thank Dr. Johan de Rooij (Utrecht University, NL) for kindly providing cells and Dr. Hedde van Hoorn (VU University, NL) for his assistance with pillar deflection analysis. Financial support for this work came from the Netherlands Organization for Scientific Research (FOM 09MMC03).

## DATA AVAILABILITY

All data are available on request to either of the corresponding authors.

## Supplemental Material

### Obtaining the cdf

For N localizations, the total number of distances between localizations is given by N^2^-N. For a typical dSTORM experiment with 10^6^ localizations this would mean 10^12^ distances. If stored as double precision values this would require 8 TB of memory, well beyond the limits of modern day PC’s. Therefore our algorithm only takes distances into account that are smaller than a set value r_max_. When 10^6^ distances are found it terminates.

### Relation between variance and squared mean

The second factor in eq.(11), characterizes the joined statistics of the photophysics of the fluorophore and the statistics of labeling of the primary antibody by the secondary antibodies.

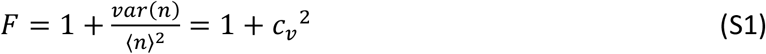

The factor relates to the coefficient-of-variation, c_v_ of n = σ_n_/<n>, in statistics. Values for F vary between 1 and 2 depending on the underlying and dominant statistics.

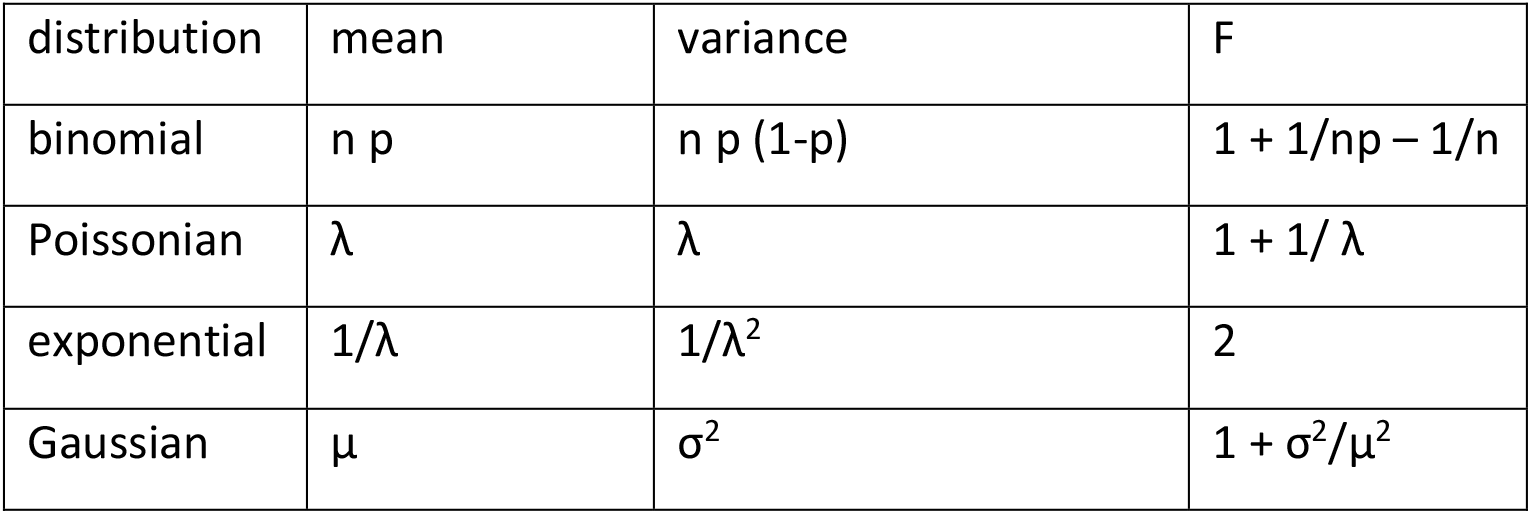

| distribution | mean | variance | F |
| --- | --- | --- | --- |
| binomial | $n p$ | $n p (1-p)$ | $1 + 1/np - 1/n$ |
| Poissonian | $\lambda$ | $\lambda$ | $1 + 1/\lambda$ |
| exponential | $1/\lambda$ | $1/\lambda^2$ | 2 |
| Gaussian | $\mu$ | $\sigma^2$ | $1 + \sigma^2/\mu^2$ |

For all, excluding the exponential distribution, F approaches 1 for large enough n. Exponential statistics would become dominant when single fluorophores are considered. In that case F = 2.

### Simulation for a combined statistics with secondary antibody labeling

A typical dSTORM experiment involves a dual labeling step where the molecule of interest is first labeled by a specific primary antibody that is subsequently labeled by multiple secondary antibodies, each conjugated to multiple fluorophores. To assess the distribution in this experiment we performed simulations. In those simulations we assumed:

1. The number of secondary antibodies bound to a primary antibody is constant, given that the excess of secondary antibody occupies all binding sites on the primary antibody.
2. The number of fluorophores bound to a secondary antibody has a Poissonian distribution with a mean of 4.7 (typical mean value provided by the manufacturer, Jackson Immunology).
3. The number of detections per fluorophore follows a single-exponential distribution, typical for photobleaching. The number of detections when a fluorophore is in the on-state equals, t_on_ × framerate. Alexa647, as used in the current study is generally assumed to behave according to a four-state molecule characterized by a ground, a fluorescent excited, a non-fluorescent triplet and a long-lived dark state. The latter is populated via the excited triplet state ^1^. The distribution in such a case is described in terms of a static trap model ^2^, with on-times following a single exponential distribution.

Figure S4 summarizes the result of this simulation. The factor 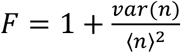, is dominated by the number of secondary antibodies. For typical values found in literature for the secondary to primary ratio (4), F is found to be below 1.1. Even in the case of only a single secondary per primary F equals 1.5, which is still below its maximal value of 2. This is caused by the multiple fluorophores per secondary antibody.

**Figure S4:**
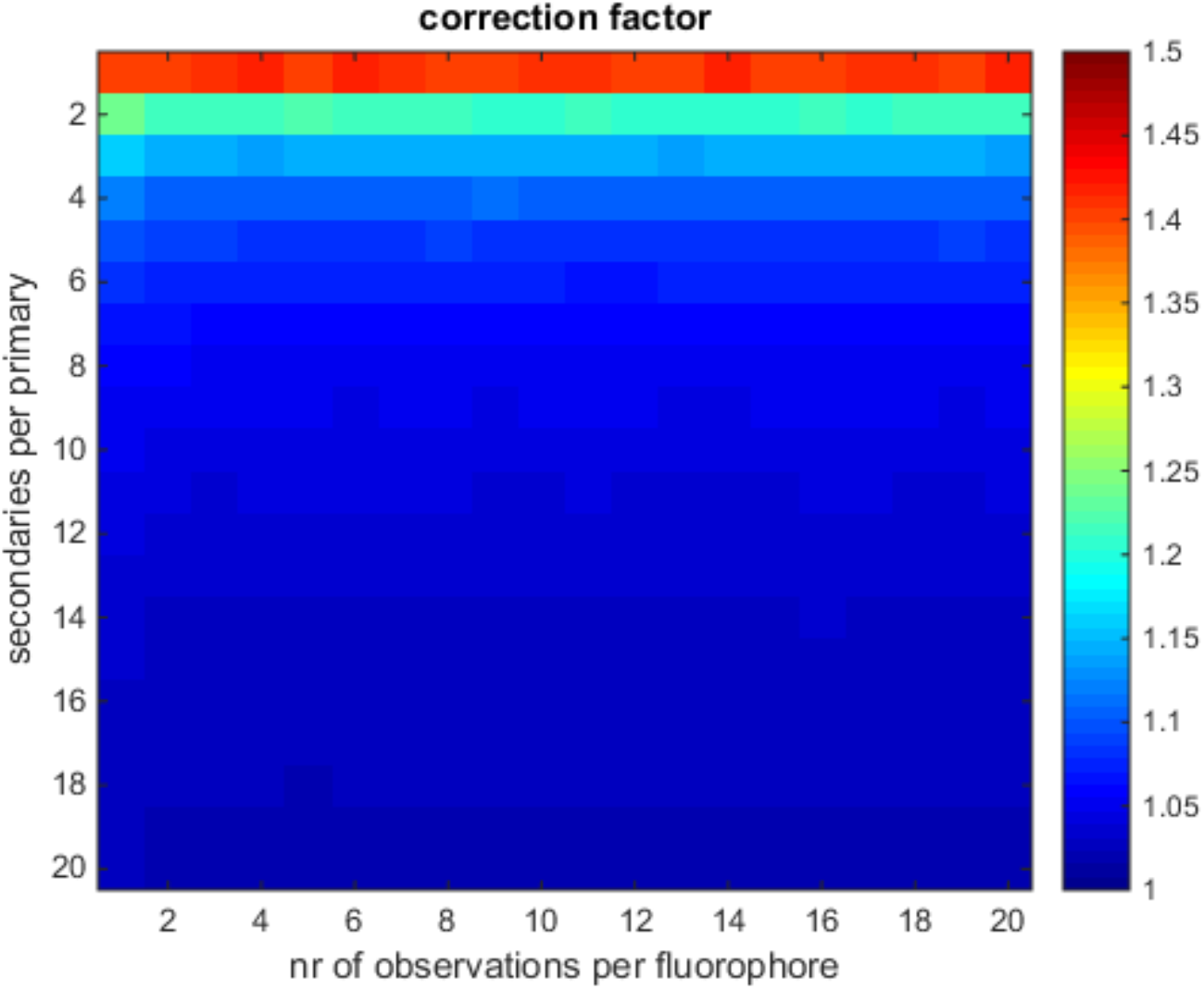
Correction factor F as described in formula 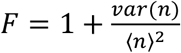 for several simulations where the number of observations per fluorophore and the number of secondary antibodies per primary antibody were varied. The number of secondary antibodies per primary dominates the factor F.

## Supplementary Figure Legends

**Supplementary Figure 1:**
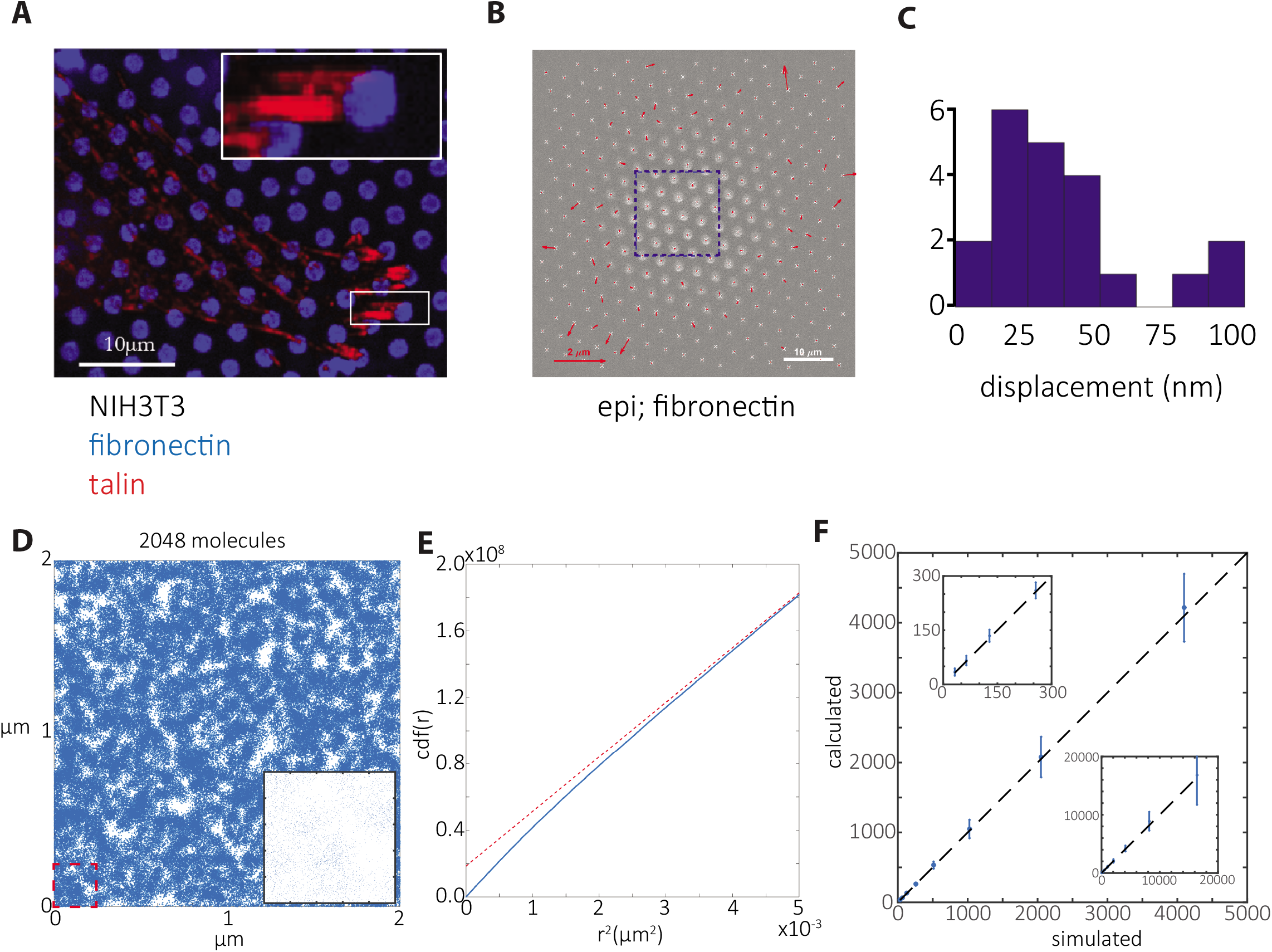
Pillar localization precision and analyses on simulations of molecules. **A,** confocal image of a NIH3T3 cell on fibronectin (conjugated with Alexa 405) stamped PDMS pillars immunostained for talin (secondary antibody Alexa 647). **B,** epi-fluorescence image of cell-free (force-free) PDMS pillar array obtained from dSTORM microscope with calculated pillar deflections (1024×1024 pixels). Blue box indicates the area in which dSTORM measurements were performed (256×256 pixels). **C,** the histogram of pillar deflections measured in the boxed area (B) indicating the localization precision of pillar centers. **D-E,** detections from a simulation of random placement of 2048 molecules in a region of 4 μm^2^ with 100 localizations per molecule (N=2048×100) with a positional accuracy of 20 nm (D), and cumulative distance function (cdf) of the inter-localization distances with a linear fit (red dashed line) with y-intercept at N_c_=2×10^7^, resulting in calculated N_m_=N^2^/(N+N_c_)=2076 (E). **F,** calculated number of molecules Nm with standard deviation plotted against simulated number of molecules and dashed line of slope 1 with insets showing same graph with different zoom areas. Scale bars are 10 μm (A, B); deflection arrow scale is 2 μm (B).

**Supplementary Figure 2:**
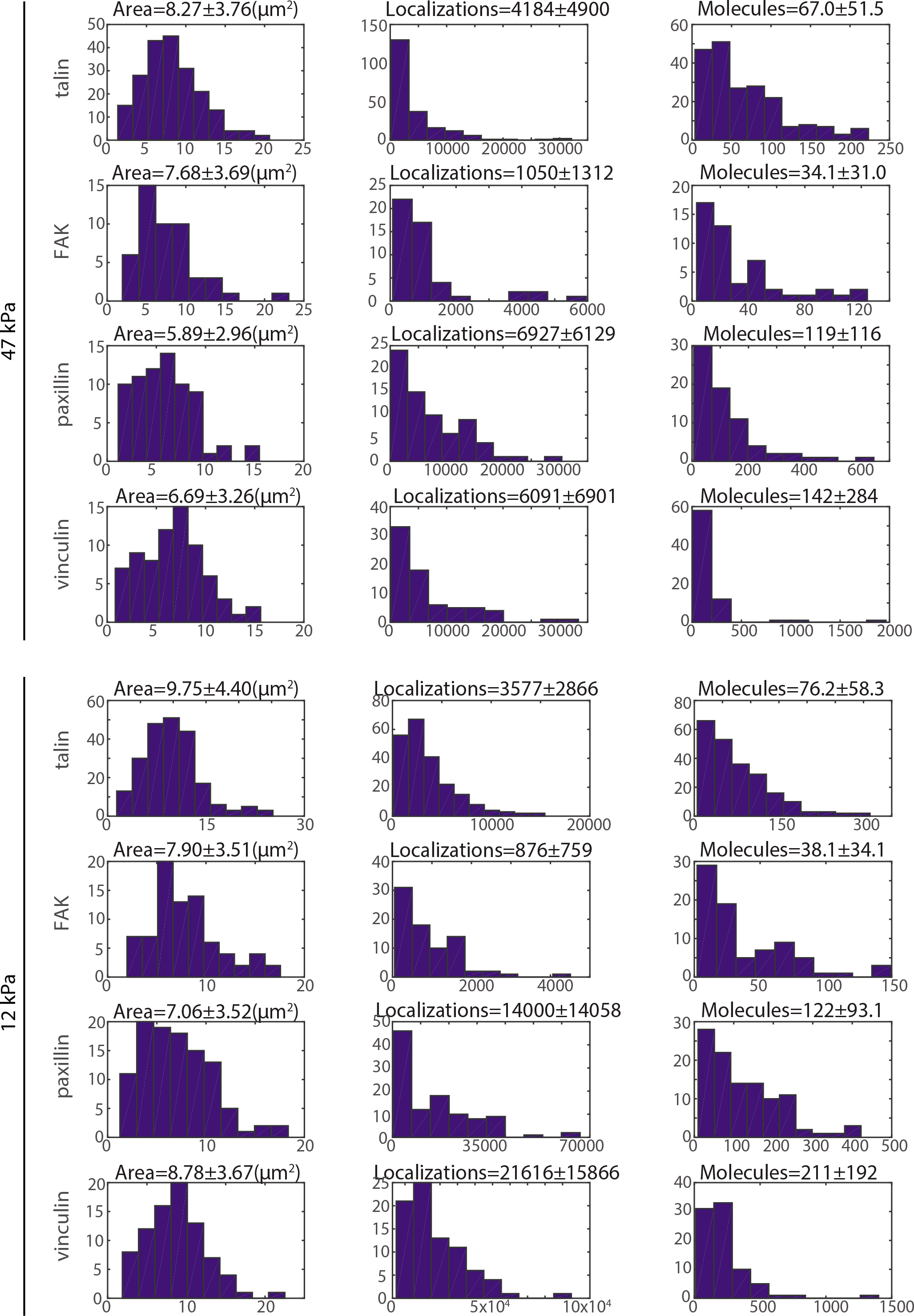
Histograms of adhesion area, localizations detected and molecules calculated per adhesion. Histograms of manually selected cell matrix adhesions associated with pillars showing distribution of total cell matrix adhesion area (first column), detected localizations (second column), and calculated number of molecules (third column). Top four rows show data for adhesions coupled to pillars with effective Young’s modulus of 47 kPa. Bottom four rows show data for adhesions coupled to pillars with effective Young’s modulus of 12 kPa. Data for dSTORM experiments on talin (1^st^ and 5^th^ rows), FAK (2^nd^ and 6^th^ rows), paxillin (3^rd^ and 7^th^ rows) and vinculin (4^th^ and 8^th^ rows) are shown. Means and standard deviations are given above each histogram.

**Supplementary Figure 3:**
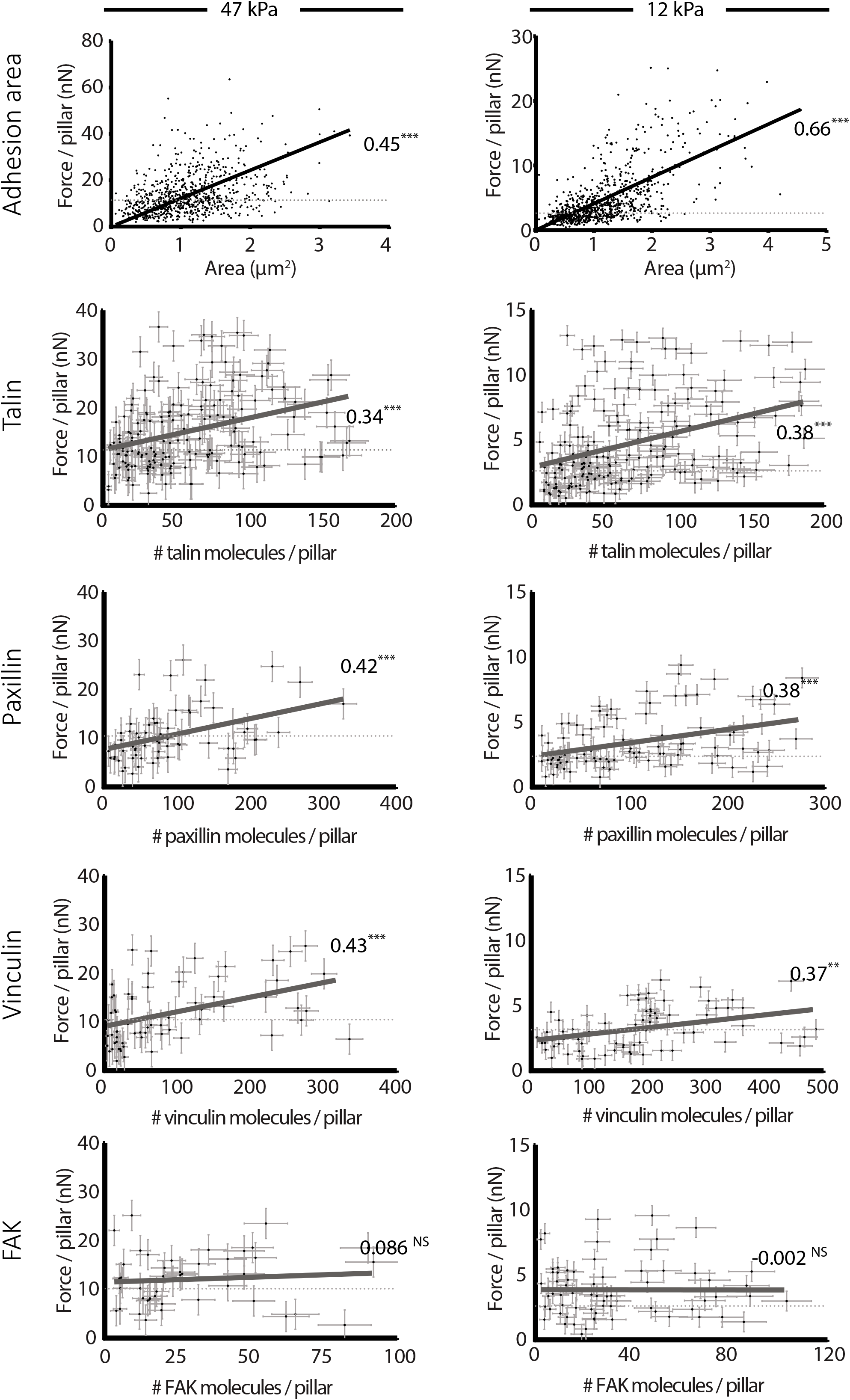
Total adhesion area and number of talin, vinculin, and paxillin molecules but not of FAK molecules correlates with local traction force. Force per adhesion measured plotted against adhesion area and number of calculated talin, paxillin, vinculin and FAK molecules associated with the adhesion from three different experiments plotted with standard deviations; black solid line is the accompanying linear fit denoted with the calculated Pearson’s correlation and dashed line is the measured background deflections for cells seeded on substrates with effective Young’s modulus 47 kPa (left) and 12 kPa (right). ***, p<0.0001; **, p<0.005; NS: p>0.05: p values denote how significantly the slope is different from zero as calculated with F-test.

